# Grammars of action in human behavior and evolution

**DOI:** 10.1101/281543

**Authors:** Dietrich Stout, Thierry Chaminade, Andreas Thomik, Jan Apel, Aldo Faisal

## Abstract

Distinctive human behaviors from tool-making to language are thought to rely on a uniquely evolved capacity for hierarchical action sequencing. Unfortunately, testing of this idea has been hampered by a lack of objective, generalizable methods for measuring the structural complexity of real-world behaviors. Here we present a data-driven approach for quantifying hierarchical structure by extracting action grammars from basic ethograms. We apply this method to the evolutionarily-relevant behavior of stone tool-making by comparing sequences from the experimental replication of ˜2.5 Mya Oldowan vs. more recent ˜0.5 Mya Achuelean tools. Results show that, while using the same “alphabet” of elementary actions, Acheulean sequences are structurally more complex. Beyond its specific evolutionary implications, this finding illustrates the broader applicability of our method to investigate the structure of naturalistic human behaviors and cognition. We demonstrate one application by using our complexity measures to re-analyze data from an fMRI study of tool-making action observation.

## Introduction

For more than 60 years, the serial ordering of behaviour has been a core topic for the cognitive and behavioral sciences^1,2^. Enhanced capacities for complex action sequencing support distinctive human behaviors such as language^3^, imitation^4^ and tool-use^5,6^, and are fundamental to the flexibility that is a hallmark of human intelligence ^7,8^. It has been suggested that this implies a unitary evolutionary and neural foundation for human cognitive uniqueness across domains^1,5,6^, but this remains controversial^9^. Although modelling suggests computational similarities across behaviours ranging from foraging to language-learning^10^ empirical investigation has been limited by a lack of objective, generalizable methods for describing, quantifying, and comparing the sequential structure of diverse, real-world behaviours. In Paleolithic archaeology, for example, investigation of long-standing hypotheses about the evolutionary relationships between tool-making, language, and cognition have been hampered by the lack of an objective metric for the behavioural complexity of different ancient technologies^11–13^. Here we adopt a data-driven computational approach to this challenge by using grammatical pattern recognition algorithms to measure the structural complexity of behavioral sequences from modern tool-making replication experiments – effectively extracting action grammars for critical survival skills from the human evolutionary past.

We conducted 17 tool-making replication experiments and coded the behavior sequences that were generated (Fig. 1A). This sample includes 5 sequences for which upper limb movements and manual joint angles were recorded as part of a previous study^14^, and 6 for which the tools and waste produced were analyzed and compared with actual Paleolithic artifacts from the Middle Pleistocene site of Boxgrove, UK^15^. Building on this and other previous research ^14–19^, we focused our current study on archaeologically documented tool-making methods from the early and late Lower Paleolithic, a period that witnessed a nearly 3-fold increase in hominin brain size. This allows us to empirically address the over 100 years of theorizing linking increasingly complex tool-making with brain evolution and language origins ^5,20–22^. The early (Oldowan, ca. 2.5 Mya) technology modeled here comprised the production of simple, sharp-edged stone flakes by striking one stone with another. The late (Late Acheulean, ca. 0.5 Mya) technology comprised the production of refined, teardrop-shaped “handaxes” through intentional shaping. We defined a shared action alphabet, consisting of 7 event types encompassing the elementary body movements and object transformations present in every sequence of both technologies, and applied two pattern recognition algorithms to the coded event sequences: Hidden Markov Modeling (HMM) and k-Sequitur.

**Fig. 1.**
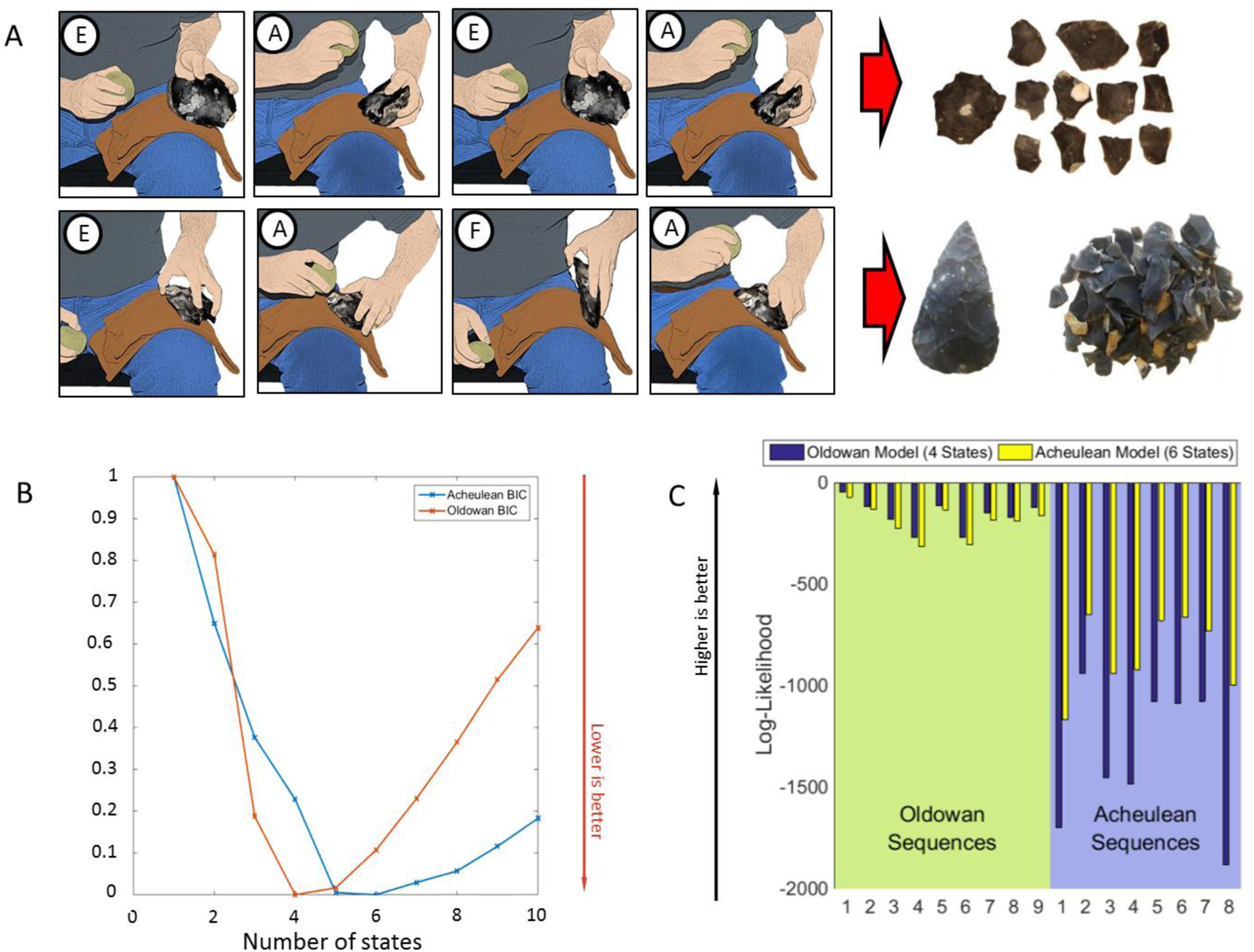
Hidden Markov Modeling of tool-making action sequences. **(A)** Oldowan (top) and Acheulean (bottom) action sequences were coded using 7 event codes (circled letters, see Materials & Methods). Products illustrated to the right. **(B)** Bayesian Information Criterion values (less is better) across models with increasing numbers of hidden states. Red, Oldowan, Minimum =4; Blue, Acheulean, Minimum=6. **(C)** Log-Likelihood values indicating model fit (higher is better) across sequences. Fit for Oldowan sequences is better overall; Acheulean model fit to Oldowan data is better than Oldowan model fit to Acheulean data.

## Results

### Hidden Markov Modeling

HMM detects probabilistic regularities (hidden states) across sequences and can capture the structure of arbitrarily complex sequences given sufficient numbers of hidden states. The optimal number of hidden states provides a measure of structural complexity. We fitted HMMs to coded event sequences, and computed the Bayesian Information Criterion (BIC) across different numbers of hidden states as a measure of model fit. BIC reached its minimum (less is better) at 4 hidden states for Oldowan and 6 for Acheulean data (Fig 1B), indicating a 50% increase in complexity. These two models perfectly categorized the sequences (likelihood greater for correct model, Fig. 1C). The fit was better for both models on the simpler Oldowan sequences. The close fit of the Acheulean model to Oldowan data (but not vice versa) indicates that the former captures most of the structure of the latter, and that Oldowan sequences may be considered a subset of Acheulean sequences.

We therefore used the Acheulean HMM to test for further structure. We obtained the most likely hidden state sequences for the Oldowan and Acheulean data and then fitted a second, 2-state HMM onto these higher-order sequences (Fig. 2). We found that Acheulean sequences oscillate between two superordinate states-of-states (SoS) whereas Oldowan sequences remain in one). Thus, Acheulean sequences display an additional level of structure not expressed by Oldowan sequences.

**Fig. 2.**
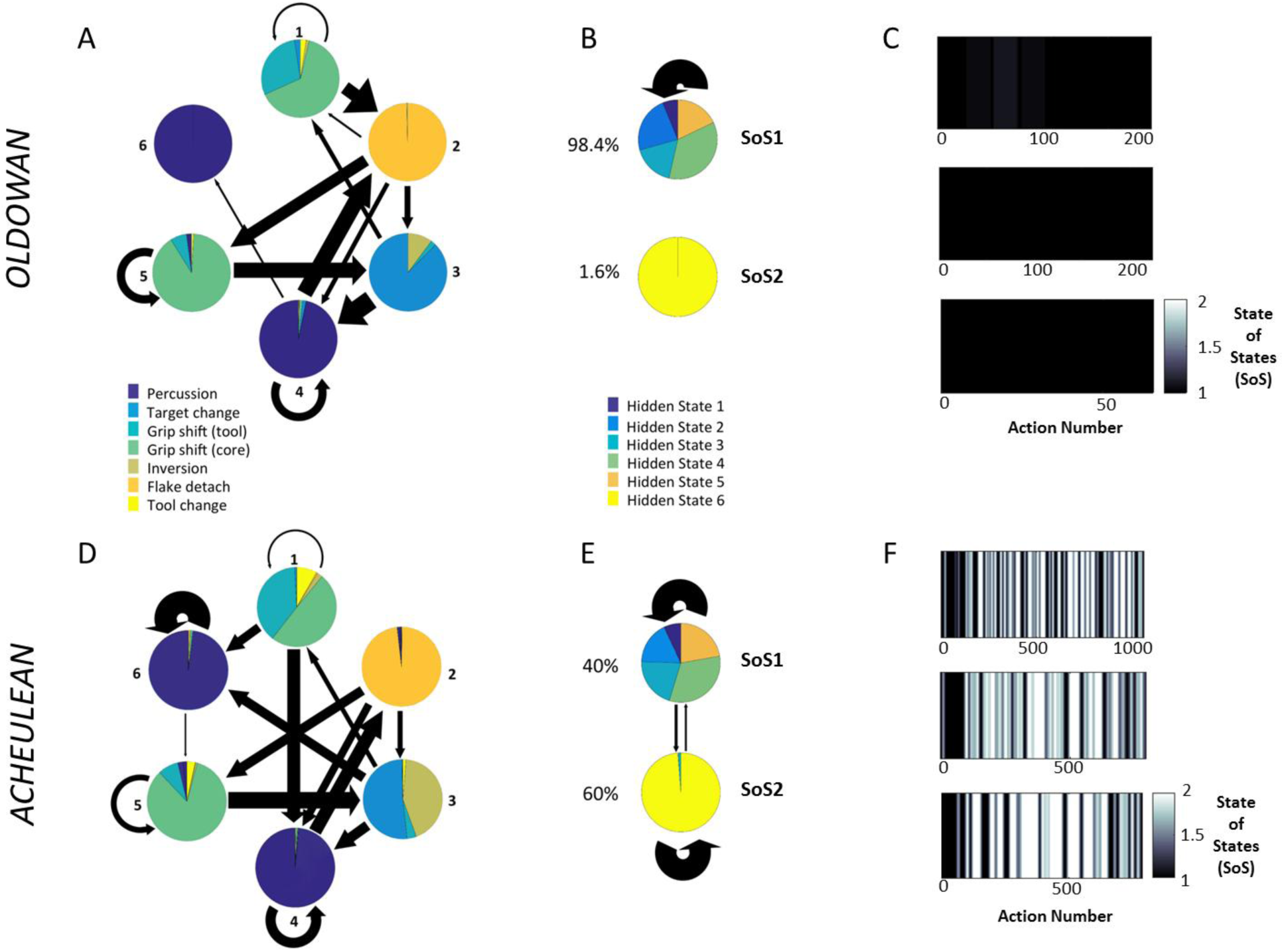
Empirical transition (arrows) and emission (pie charts) matrices of 6-state HMM fitted to all Oldowan (A) and Acheulean (D) sequences. Arrow thickness indicates transition probabilities between states (values <5% not displayed). Pie chart area indicates probability of an action being performed in that state. Oldowan Hidden State 6 accounts for less than 2% of all data points. In the middle are similar illustrations of the superordinate “States-of-States” for Oldowan **(B)** and Acheulean **(E)** data. At right are examples of the running average State of States for Oldowan **(C)** and Acheulean **(D)** time-series. Black: everything in SoS 1; white: everything in SoS 2.

Next, we fit the 6-state Acheulean HMM to Oldowan and Acheulean data and observed the probability of actions per hidden state as well as transitions between hidden states. Our Oldowan data are characterized by the repetition of one simple action “chunk” consisting a relatively invariant sequence of states (3->4->2: Fig. 2A) that essentially corresponds to the removal of an individual flake and is entirely captured by SoS1 (Fig. 2B&C). Acheulean sequences are more variable (Fig. 2D), reflecting the addition within some flake removal chunks of a sub-operation archaeologists refer to as striking platform preparation. This involves repeated low-amplitude (see Methods and Supplementary Fig. 1) chipping of striking surfaces to alter their sharpness, bevel, and placement relative to the midline. This operation is captured at the next level by SoS2 (Fig. 2E). SoS2 is less frequent in the early stages of our sequences (Fig. 2F) which is consistent with the presence of an initial “roughing out” stage in handaxe manufacture prior to more refined shaping ^23^. Introspection by experienced tool-makers^12,17,24^ has previously suggested that platform preparation increases the complexity to Paleolithic action organization, but it has not previously been possible to test this intuition objectively or to quantify the magnitude of increase in a generalizable way. Our HMM method thus captures meaningful (i.e. goal directed) regularities in stone tool-making in a data-driven way that: 1) derives structure rather than imposing it, 2) respects the real variability underlying ideal characterizations, 3) enables objective quantification of grammatical complexity, and 4) is readily adaptable to the study of other sequential behaviors.

The Chomsky hierarchy in Formal Language Theory (FLT) describes a series of increasingly powerful and inclusive computational systems, or grammars, differentiated by their memory resources ^1,25^. A simple Markov chain is a memoryless probabilistic system equating to a regular (finite-state) grammar that does not permit long distance dependencies. HMMs are dynamic Bayesian networks that asymptotically approximate supra-regular context-free grammars (with unbounded memory) through the progressive addition of hidden states. The increase we observed in the optimal number of hidden states from Oldowan to Acheulean thus provides a measure of increased grammatical complexity and memory requirements without positing infinite capacity. This modeling approach is consonant with the view that finite-state, probabilistic, and parallel computational models are cognitively and neurobiologically realistic ^7,25–27^. Others, however, contend that human cognition is in fact characterized by constitutively hierarchal processing using supra-regular resources and that humans have a tendency to employ such context-free solutions even when they are not actually necessary ^28,29^.

### Context-free grammar fitting

We therefore pursued a second approach by fitting context-free grammars (CFGs) to the tool-making sequences. FLT employs terminal symbols (in our case 7 event types) and non-terminal symbols (re-write rules expandable to terminal and/or non-terminal symbols) to generate strings. Whereas regular grammars and HMMs are driven by local relationships between symbols, CFGs capture nested dependencies of theoretically infinite length and depth. The standard algorithm to extract CFGs, Sequitur ^30^, creates a new rule as soon as a symbol pair is observed twice in a sequence and repeats this pair-wise aggregation, adding new levels of superordinate rules until the complete sequence is described. This makes Sequitur powerful but liable to detect a high number of spurious (occurring <3 times) rules in the variable sequences generated by real human behavior. We therefore developed an algorithm, k-Sequitur, requiring a pair to occur k-times before generating a rule. Increasing k makes the grammar discovery process less sensitive to infrequent pairs and less prone to creating rules from noise.

In agreement with our HMM results, CFG extraction found Oldowan grammars to be a less complex sub-set of Acheulean grammars. Rule inference from combined Oldowan and Acheulean samples identified multiple rules that occur only in Acheulean sequences (Supplementary Fig. 2) and showed that the frequency of Acheulean-only rules increases at higher levels (0 at level 2, 1 at Level 3, 2 at Level 4, 5 at Level 5). No Oldowan-only rules were identified, even when rule inference was restricted to the Oldowan data set. CFG extraction achieved substantial compression of both Oldowan and Acheulean sequences (Fig. 3A), however the rate (inverse slope) of Acheulean compression was more than twice as great (7.69 vs. 2.94). This indicates that Acheulean sequences have more structure for rule-based compression, in a ˜2:1 ratio paralleling our HMM finding of two Acheulean SoS vs. one Oldowan. Each post-compression Acheulean element (rule or terminal symbol) contains more information (measured as Shannon entropy: Fig. 3B), yet Acheulean grammars still require more non-terminal symbols (rules) to achieve a comparable fit to the data. These compression results are robust over increasing k values (Supplementary Fig. 3). CFGs can parse regular strings, so fitting CFGs to our sequences in this way does not imply that supra-regular resources are required. It does show that the greater complexity and depth of Acheulean sequences is robust even assuming such resources.

**Fig. 3.**
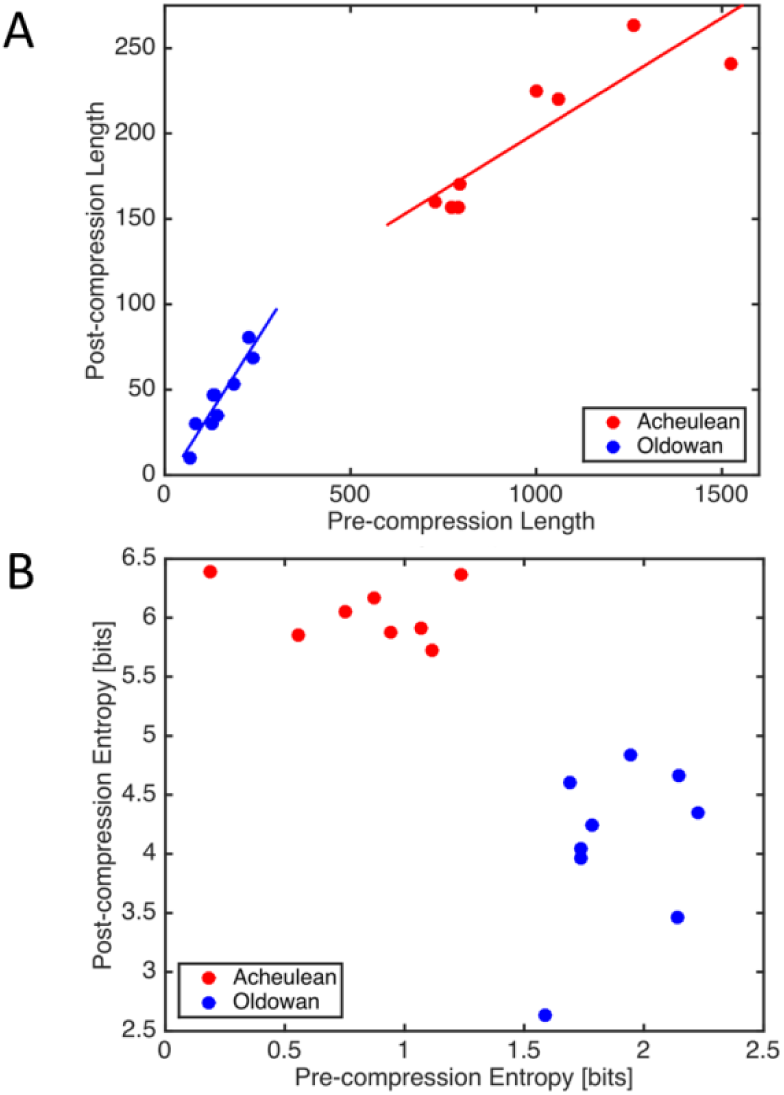
Effect of Sequitur compression on sequence length (A) and entropy (B).

Further inspection of CFG results reveals that the greater complexity of Acheulean sequences is due to long strings of repeated percussions, the removal of which eliminates Oldowan/Acheulean differences in compression rate (Supplementary Fig. 4). These strings comprise the same repeated, low-amplitude chipping of striking platforms (Supplementary Fig. 1) extracted as SoS2 in our HMM analysis and corresponding to the tool-making operation known as platform preparation^15^. HMM and CFG methods thus converge, not only to quantify the greater complexity of Acheulean sequences, but also to extract a key technological element of the instrumental structure of Acheulean tool-making that largely accounts for this difference.

**Fig. 4.**
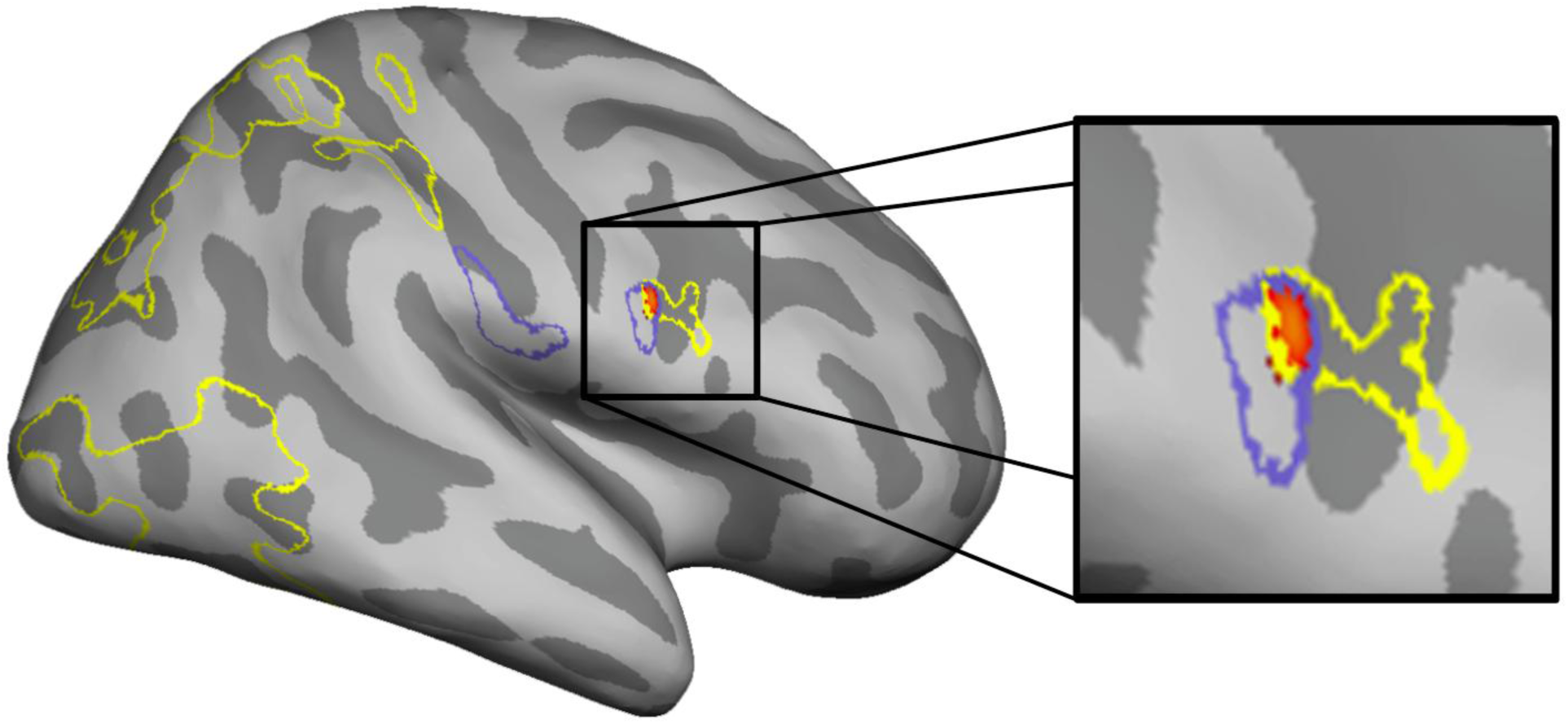
Covariance of BOLD response with tool-making stimulus behavioral complexity. Outlines corresponding to areas yielding significant negative correlation with the CFG (purple) and HMM (yellow) covariates describing action sequence complexity. Heatmap clusters represent the minimum of these two correlations where they overlap voxel-wise in the right pars opercularis of the inferior frontal gyrus (right).

## Discussion

Results indicate that our grammar extraction methods are able to discover the instrumental structure of behavior directly from the correlational structure of action sequences coded using a minimalistic and objective ethogram, without requiring subjective functional or intentional interpretations by the observer. These methods are easily generalizable to other behaviors and, in the specific case of Paleolithic tool-making, provide new means to investigate the archaeological record of technological change and variation. By using a single elementary action alphabet it is possible to consider variation within as well as between different types of tool-making in strictly equivalent terms, treating behavioral variation as a source of information rather than noise and avoiding problematic assumptions regarding the behavioral reality these archaeologically-constructed types.

Our specific findings focus attention on the emergence of platform preparation techniques during later Acheulean times^15^ as one key indicator of increasing technological complexity. Modern experimental tool-makers have identified platform preparation as an important technique enabling production of the large, relatively thin flakes needed to thin (“refine”) a handaxe without a disproportionate decrease in breadth^31^. Behavioral variation across modern knappers at different levels of expertise reveals that platform preparation is a difficult technique to master in application^15^, even if its function is clearly understood. Expert intuitions regarding the behavioral and cognitive complexity ^12,17,24^ of platform preparation have motivated the proposal that its emergence and spread in Late Acheulean (˜780 - 400,000 years ago) times may be related to the rapid encephalization that also occurred during this time^15^ (see Supplementary Discussion).

Neuroimaging results have consistently linked Late Acheulean style handaxe-making with the right Inferior Frontal Gyrus (rIFG). This includes evidence of increased functional activation during action execution^17,32^ and observation^18^, as well as structural remodeling of underlying white matter in response to tool-making training^19^. To date, these findings have been interpreted using reverse inference from known functions of rIFG, including especially its involvement in the cognitive control functions of inhibition^33^ and action updating^34^ that are critical to response selection during multi-component behavior^35^. Direct manipulation through transcranial magnetic stimulation confirms the causal role of rIFG in such selection^36^. We have argued^19,37^ that this domain-general computational function unifies superficially diverse evidence of rIFG involvement in tasks ranging from fine motor control and manual sequence learning to linguistic processing of syntactic violations and perturbations. Combined with evidence that rIFG connections to parietal cortex via the 3^rd^ branch of the Superior Longitudinal Fasciculus (SLFIII) are evolutionarily expanded in humans^37^ and plastically remodeled by tool-making training^19^, this interpretation provides functional, archaeological, and neuroanatomical grounding for longstanding hypotheses of tool-language co-evolution^5,6,20,21,38,39^. However, such reverse inference from observed activation to implied mental process is potentially problematic given the likelihood that any particular brain region might support multiple functions^40^. This is ameliorated to some extent by defining “function” in terms of underlying computational roles rather than specific behavioral tasks, but nevertheless remains an issue. One strategy to address this concern, is to strengthen the basis for reverse inference by more rigorously defining the task context^41^ although this can be quite difficult for naturalistic behaviors like stone tool-making.

In fact, neuroimaging studies of stone tool-making^17,18,32^, have largely been limited to categorical comparisons between abstracted tool-making types that do not dissect the contribution of specific behaviors, like platform preparation, that are variably involved across particular instances. Artificial tasks designed to isolate such elements of stone tool-making have provided insight into the demands of specific behavioral components^42,43^, but this approach sacrifices ecological validity and cannot address real-world behaviors that necessarily differ on multiple dimensions at the same time. For example, similar challenges confront attempts to study language as it is actually used by people to communicate rather than in the form of controlled laboratory manipulations^44–46^. In the current case, it is not possible to control lower-level kinematic, geometric, and visual features of stone tool-making without also altering the higher-level action structure that emerges from them and seriously compromising the basis for analogy with real, archaeologically-documented behaviors. An alternative approach, developed here, is to derive behavioral models which make specific neuro-cognitive processing predictions that are testable against brain data^46^.

To illustrate this approach, we used HMM and CFG grammar extraction to measure the complexity of action sequences in Oldowan and Acheulean video stimuli from a published fMRI study of tool-making action observation^18^. In common with tool-makers in the current study, the demonstrator recorded in these stimuli was instructed to emulate handaxes from the Acheulean site of Boxgrove. Comparative artifact analyses have confirmed that tool-making methods in both studies closely approximate actual Paleolithic behavior at Boxgrove, specifically including the use of platform preparation^15^. To generate a continuous complexity measure from HMM, we used the difference in likelihood (measured by Akaike Information Criterion, see methods) between more (6 state) and less (4 state) complex models fit to the stimulus sequences. For CFG, we simply used the compression ratio. In both cases, increasing complexity is indicated by lower values, so we expected a negative correlation with BOLD response in subjects attempting to parse the structure of observed sequences.

Results (Fig. 4) suggest that HMM and CFG capture partially overlapping stimulus processing demands. Interestingly, differences appear related to sensory modality. Increasing HMM complexity uniquely recruits occipitotemporal and parietal portions of a dorsal attention network ^47^ responsible for the intentional allocation of visual attention whereas unique CFG response occurs in somatosensory cortex of the right parietal operculum, plausibly reflecting vicarious tactile processing ^48^. An interesting direction for future work might be to investigate why these different computational approaches appear to map to different sensory modalities.

The conjunction of the two covariates reveals regions of left parietal operculum and rIFG (*pars opercularis*) that are specifically responsive to stimulus complexity irrespective of measurement method. This is generally consistent with the well-documented involvement of ventral frontoparietal cortex in naturalistic stone tool-making^17,18,49^ and specifically supportive of the functional interpretation of rIFG response to handaxe production presented above. Importantly, this finding is independent of experimental condition (Oldowan, Acheulean) and thus strengthens the inference that it is indeed continuous variation in behavior complexity, rather than some other confounding difference between stimulus types, that is driving the response. To further strengthen this inference, future research should combine grammar extraction methods presented here with kinematic recording and eye-tracking to simultaneously explore brain response to multiple dimensions of behavioral variation. Such methods have already been applied to demonstrate that experimental Oldowan and Acheulean tool-making (including 5 sequences used here for grammar extraction) do not differ in manipulative complexity as measured by the statistics of manual joint movements^14^.

The remarkable expressive power of human language derives from an ability to recombine a relatively small set of discrete units into a vast array of meaningful structures ^3,29^. At the outset of the Cognitive Revolution, Lashley ^2^ used the example of language to argue that *all* skilled behavior (and associated neural activity) is organized in this hierarchical fashion. This insight that was subsequently applied to the specific case of stone tool-making by Holloway^5^. Sixty-odd years later, however, we are still struggling with what Lashley (p. 122) identified as the “essential problem of serial order”: defining the “generalized schemata of action which determine the sequence of specific acts” that he termed “the syntax of the act.” Here we developed objective and generalizable methods for defining and quantifying these structures (action grammars) along with their neural correlates from raw behavioral data. While the detailed kinematics of hand actions to produce a tool vary from trial to trial considerably, we found an invariant hierarchical structure underpinning performance. Our analytical approach does not postulate the existence of such “action grammars” *a priori*, but instead identifies them from raw behavioral data using machine learning techniques, showing that even with the same alphabet of actions qualitatively more complex artefacts can be produced by using measurably more complex action grammars. In addition to the method’s broad utility for the behavioral and social sciences, the finding that our automatic identification of action grammars maps to distinct neural correlates offers the potential for novel quantitative approaches to the hierarchical structure of behavior across applications from dexterous prosthetics ^50^, to the training of surgeons ^51^ and human-like AI ^52^.

## Methods

### Tool Replication

Tool replication was performed by two expert stone tool-makers (knappers) with decades of experience. The research was approved by the University College London Research Ethics Committee [0603/001] and each participant provided written informed consent. All experiments were video-recorded, including 9 instances of Oldowan knapping and 8 instances of Acheulean knapping. In each experiment, a piece of flint was worked until either completely exhausted (Oldowan) or successfully shaped into a refined handaxe (Acheulean). Six of the handaxes produced in these experiments have previously been described and compared to archaeological examples for the Middle Pleistocene site of Boxgrove ^15^. Kinematics from a different subset of the experiments (3 Oldowan, 2 Acheulean) have been published ^14^.

Experimental replication is a long-established research method in archaeology, especially with respect to flaked stone technology ^53^. Our tool-making experiments drew upon this background to model simple flake production (cf. “Oldowan”, “Mode 1”, “Mode C”^54^, here termed “Oldowan”) and refined handaxe shaping (cf. “Later Acheulean handaxe”, “Mode 2”, “Mode E2” ^54^, here termed “Acheulean”). Previous experimentation has shown that a wide range of Oldowan forms may be replicated through hard-hammer free-hand percussion without intentional core shaping ^55^, whereas other techniques (e.g. bipolar, passive hammer) produce diagnostic traces that are less common in the archaeological record ^56^. Although there is some evidence of structured reduction strategies in the Oldowan (e.g. preference for unifacial vs. bifacial flaking ^57^) it is possible to produce most or all Oldowan forms through unstructured (cf. “mindless” ^58^ or “least effort” ^55^) flaking. We thus instructed our subjects to knap Oldowan experiments in an opportunistic fashion, following the definition of “simple debitage” provided by ^59^. For Acheulean experiments, subjects were instructed to produce “refined” Acheulean handaxes of the kind known from the site of Boxgrove (with which subjects were familiar). This included the use of soft hammers and simple platform preparation (faceting), both of which are attested in the Boxgrove archaeological collection ^15^. Experimental handaxes produced were comparable in refinement and debitage morphology to those from Boxgrove ^15^.

Paleolithic tool-making occurred over a vast time period and many millions of square miles, and encompasses substantial variation that could not be included in our experiments. The methods we did select are considered broadly representative of early and late Lower Paleolithic technology, and details of the production techniques employed match those documented in specific archaeological collections. We thus consider our training protocol to be both generally representative and specifically accurate in re-creating Paleolithic tool-making action sequences.

### Event Coding

We defined an action alphabet consisting of 7 event types encompassing the elementary body movements and object transformations present in every sequence of both technologies. Events were transcribed from video-recordings using Etholog 2.25 ^60^. Events were defined as follows:

A. Percussion: Striking core with percussor (hammerstone or antler billet).
B. Target Change: A change in the location of percussion on the core.
C. Grip Shift Core: Movement of the hand grasping the core.
D. Grip Shift Tool: Movement of the hand grasping the percussor.
E. Inversion: Flipping over the core without otherwise reorienting.
F. Flake detach: Removal of a flake (judged to be) > 20mm.
G. Tool Change: Exchange of one percussor for another.

This provides a minimalistic alphabet intentionally designed to limit the need for subjective interpretation and to avoid building prior hypotheses (e.g. ref. ^12^) into the coding scheme. In particular, any attempt to infer the intention of the knapper (e.g. identifying a flake detachment as “preparatory” or “thinning”) was avoided. Much richer description of knapping actions in terms of technological function is both possible and informative (e.g. ref ^61^), but was not in line with our aim to develop a data-driven and generalizable method. The coding scheme was developed through pilot work with the MRI stimulus videos (Table 1 in ref. ^18^) to be complete (every action on the core or percussor is coded), exclusive (no action could have two codes), and unambiguous. While the actual alphabet used here is specific to stone tool-making, this approach to coding could be generalized to any sequential behavior.

During transcription, we recorded an eighth event type, “Light Percussion” (Figure Extended Data Fig. 1), which was not subsequently employed in analysis. This event was defined as “Striking core with percussor using small amplitude arm movements not intended to detach flakes > 20mm” and was omitted because: 1) it required interpretation, 2) it did not occur in Oldowan sequences, and 3) it might be ambiguous with the Percussion event. Thus, we treated all instances of “light percussion” simply as percussion. However, this gesture – typical of a technical operation known as “striking platform preparation” – was rediscovered by our HMM and Sequitur analyses based purely on sequential structure (see SI), thus providing a validation of our minimalistic coding and data-driven approach.

### Hidden Markov Modeling

We fit Hidden Markov Models (HMM) to the action sequences using the Baum-Welch algorithm implemented in Kevin Murphy’s Bayes’ Net Toolbox. As the algorithm is very sensitive to the initial estimates of the transition and emission matrices, we fit each data set 1000 times for each number of states by randomly varying the initial condition and only picked the HMM with the highest log-likelihood. To compare HMMs with different number of hidden states with each other, we computed the Bayesian Information Criterion (BIC) which gives a measure of model fitness penalised by the number of free parameters in the model.

From the 6 state Acheulean HMM we obtained the most likely state-sequences through the Oldowan and Acheulean action sequences by using the Viterbi algorithm. To investigate whether the obtained hidden state sequence, contained more structure, we fitted a second, 2- state HMM onto the state sequences. As previously described, 1000 runs were performed to obtain the best-fitting HMM. Using the Viterbi algorithm again gives rise to a hidden state sequence within the hidden state sequences, a hidden “States of States” (SoS) sequence.

### Deterministic context-free grammar fitting

Any stochastic regular grammar can be represented by a uniquely corresponding HMM where outputs correspond to terminal symbols. Left regular stochastic grammars - because they are strictly equivalent to first order Hidden Markov Models - can only model phenomena with very short memory. Stochastic Context-Free Grammars represent a super-set of stochastic grammars which can feature long term memory and very hierarchical organization.

Sequitur ^30^ is a recursive algorithm that infers a hierarchical structure in the form of a context-free grammar from a sequence of discrete symbols. The Sequitur algorithm constructs a grammar by substituting repeating symbol digrams in the given sequence with new rules and therefore produces a concise representation of the sequence. The algorithm works by scanning a sequence of terminal symbols and building a list of all the symbol pairs which it has read. Whenever a second occurrence of a pair is discovered, the two occurrences are replaced in the sequence by a non-terminal symbol, the list of symbol pairs is adjusted to match the new sequence, and scanning continues. If a pair’s non-terminal symbol is used only in the just created symbol’s definition, the used symbol is replaced by its definition and the symbol is removed from the defined nonterminal symbols. Once the scanning has been completed, the transformed sequence can be interpreted as the top-level rule in a grammar for the original sequence. The rule definitions for the non-terminal symbols which it contains can be found in the list of symbol pairs. Those rule definitions may themselves contain additional non-terminal symbols whose rule definitions can also be read from elsewhere in the list of symbol pairs.

For example:

Input sequence: the little cat chases the mouse the little cat catches the mouse the big cat chases the little cat the little cat runs away from the big cat

Compressed sequence: r2 chases r3 r2 catches r3 r5 chases r2 r2 runs away from r5

Grammar:

– Root -> r2 chases r3 r2 catches r3 r5 chases r2 r2 runs away from r5
– r2 -> the little cat (used 4 times)
– r3 -> the mouse (used 2 times)
– r5 -> the big cat (used 2 times)

We ran Sequitur on each sequences in both the Acheulean and the Oldowan data sets and enumerated all the rules found across both data sets. After inferring rules from the combined Acheulean and Oldowan data set, we found that some rules only occurred in Acheulean sequences (Extended Data Fig. 2).

The Sequitur algorithm reduces the length of the sequences by replacing terminal symbol strings with aggregating rule strings. This compresses the sequence by reducing its redundancy. Figure 3A shows that sequences in our Oldowan and Achuelean samples share common compressible structure within samples but are distinct across samples. This is indicated by the fact that their pre and post-compression lengths are linear and have distinct slopes. Linear regression fit for Acheulean data is R2= 0.9852 with slope = 0.13; for Oldowan data R2=0.9982 and slope = 0.34. The inverse slope on this plot corresponds to the data compression rate through rule extraction.

Sequitur as a compression algorithm is loss-less, in that reverse applying the rules recovers the original sequence error free, and thus the same information is communicated by fewer symbols. This contrasts with the hidden states of the HMM that only capture probabilistically a higher order structure. A Sequitur compressed sequence must have more information per character and this gain in information density can be quantified using Shannon’s entropy measure. Shannon’s entropy is computed directly as the log probability of each symbol averaged over all symbols. A sequence with equally probable use of all symbols has the highest entropy, while a sequence using only a single symbol has an entropy of 0. Entropy thus measures how unpredictable a symbol is. We plotted the pre and post compression entropies in Figure 3B. Pre-compression entropy of Acheulean sequences is considerable lower than that of Oldowan sequences due to the much higher frequency of percussion events. However, post-compression entropy is considerably higher for Acheulean sequences than Oldowan sequence. Thus, pre-compression Acheulean elements (rules + symbols) carry less information than Oldowan elements whereas after compression the reverse is true.

### fMRI covariates

In order to generate covariates for fMRI analysis it was necessary to produce continuous measures of complexity for the 20 seconds video stimuli. For HMM, we first applied the method described above to each stimulus and then evaluated how well the stimulus was explained by the two respective (Acheulean 6 hidden state vs. Oldowan 4 hidden state) HMM models. Sequence length was both short and variable (stimuli were controlled for time rather than number of actions), so we employed the Akaike Information Criterion [AIC] which, unlike BIC, is not directly dependent on sample size in order to avoid confounding sequence length with model likelihood. Differences in AIC between models indicate the relative strength of evidence in their favor. Because our models differ in complexity, this difference provides a continuous measure of how complex (i.e. Acheulean-like vs. Oldowan-like) each short stimulus sequence is compared to models derived from our entire corpus. As a lower AIC indicates a more probable model, decreasing values for Acheulean – Oldowan AIC indicate increasing stimulus complexity and we predict a negative correlation with BOLD response measured by fMRI.

For CFG, we applied the same deterministic grammar extraction approach discussed above. However, in our main analysis, sequitur was applied separately to each sequence. To generate a CFG covariate comparable to our HMM AIC metric, it was necessary to generate a global set of rules derived over the entire corpus to which individual stimuli could be compared. We thus fitted sequitur to the complete set of all sequences in one run. This provided us with a sequitur parse using compressed rules for the entire corpus. We then broke down the compressed rules and matched them to the individual stimulus sequences and computed the basic metrics (as for the long sequences) for these matched compressed sequences. The compression ratio for each stimulus provides a straightforward measure of complexity, we used post-over pre-compression values so that our CFG metric would parallel our HMM metric in matching decreasing values with increasing complexity and predicting negative correlation with BOLD.

### fMRI Analyses

Experimental paradigm and participants were presented ref. ^18^. Briefly, 10 Naïve, 10 Trained and 5 Expert subjects observed 20-second videos of an expert demonstrator performing two tool-making methods of differing complexity and antiquity: the simple ‘Oldowan’ method documented by early tools 2.5 million years ago; and a more complex ‘Late Acheulean’ method used to produce refined tools 0.5 million years ago. In the present SPM analysis, the two categories of tool-making were defined as two conditions, and complexity scores (HMM and CFG) were added as covariates describing each stimulus in two individual subject analyses.

The effect of these covariates combined across the two categories of stimuli were entered in two multisubject analyses across the 20 non-expert participants, thresholded at p < 0.05 FDR-corrected at the cluster level (Fig. 4). Experts were omitted due to a small sample size insufficient to properly assess confounding expertise and automaticity effects^62,63^. To confirm the overlap in left parietal and right frontal cortices between the two analyses, a conjunction (“&”) was calculated between the T-maps describing the voxels yielding significant negative correlation with the two covariates. This analysis yielded two clusters, one in the left parietal operculum and one in the posterior part of the right inferior frontal gyrus corresponding to the pars opercularis according to the Anatomy toolbox ^64^.

## End Notes

**Supplementary Information** is available in the online version of the paper.

## Acknowledgements

This work was supported by the Commission of the European Communities Research Directorate-General Specific Targeted Project number 029065, “Hand to Mouth: A framework for understanding the archaeological and fossil records of human cognitive evolution.” We thank Francois Belletti for software development support, and James Steele and Daniel Wolpert for support and encouragement that made this project possible.

## Author Contributions

DS conceived the study and conducted the replication experiments. TC analyzed brain imaging data. AF and AT developed and applied pattern recognition methods. JA contributed to ethogram development and coded videos. DS, AF and TC wrote the paper.

## Author Information

The authors declare no competing financial interests. Data and descriptors will be published on the FigShare community archive doi: 10.6084/m9.figshare.3481958 (embargoed until publication).

## Supplementary Information

**Supplementary Figure 1.**
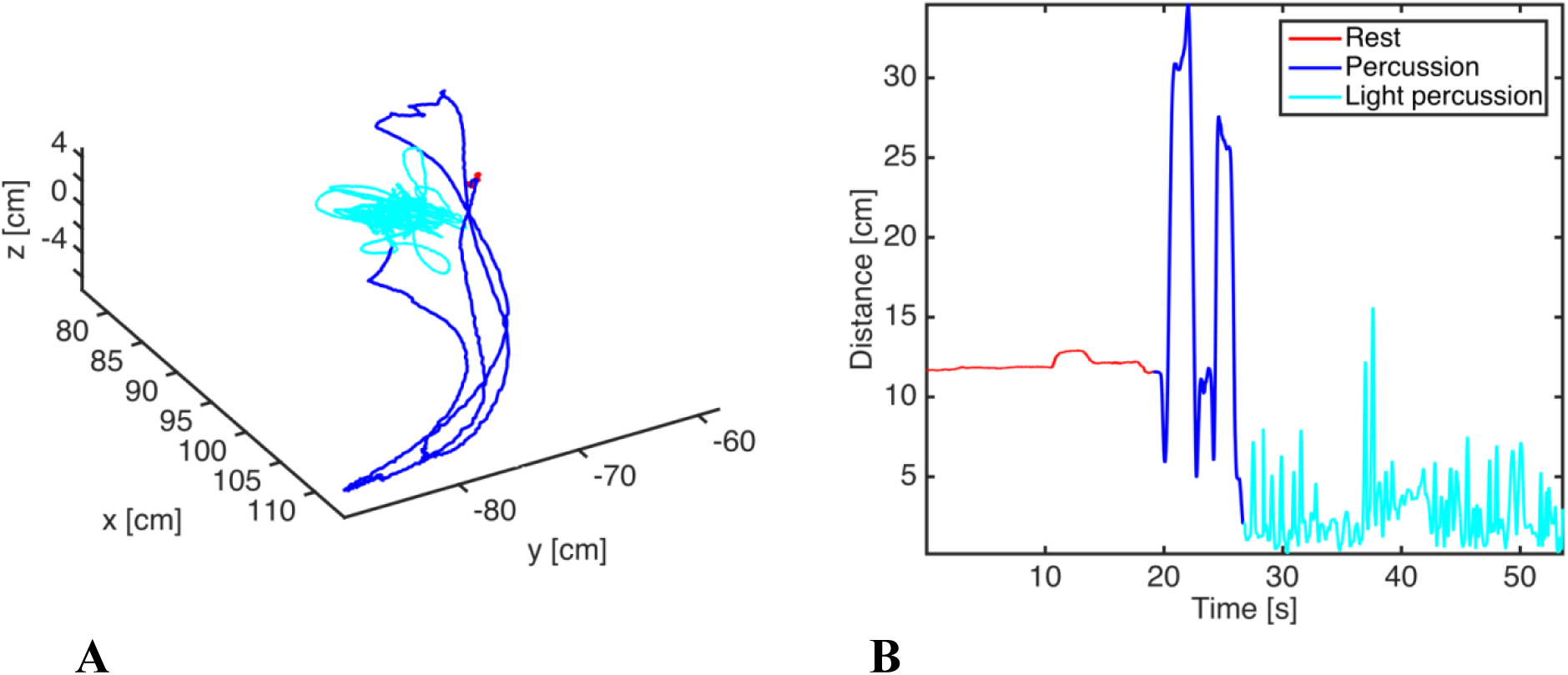
The light percussion involved in striking platform preparation (light blue) involves many rapid, low amplitude blows **(B)** when compared to more forceful percussion aimed at flake detachment (dark blue). Light percussion is achieved with wrist movements that keep the hand close to the core compared to the elbow swings deployed in forceful percussion **(A)**.

**Supplementary Figure 2.**
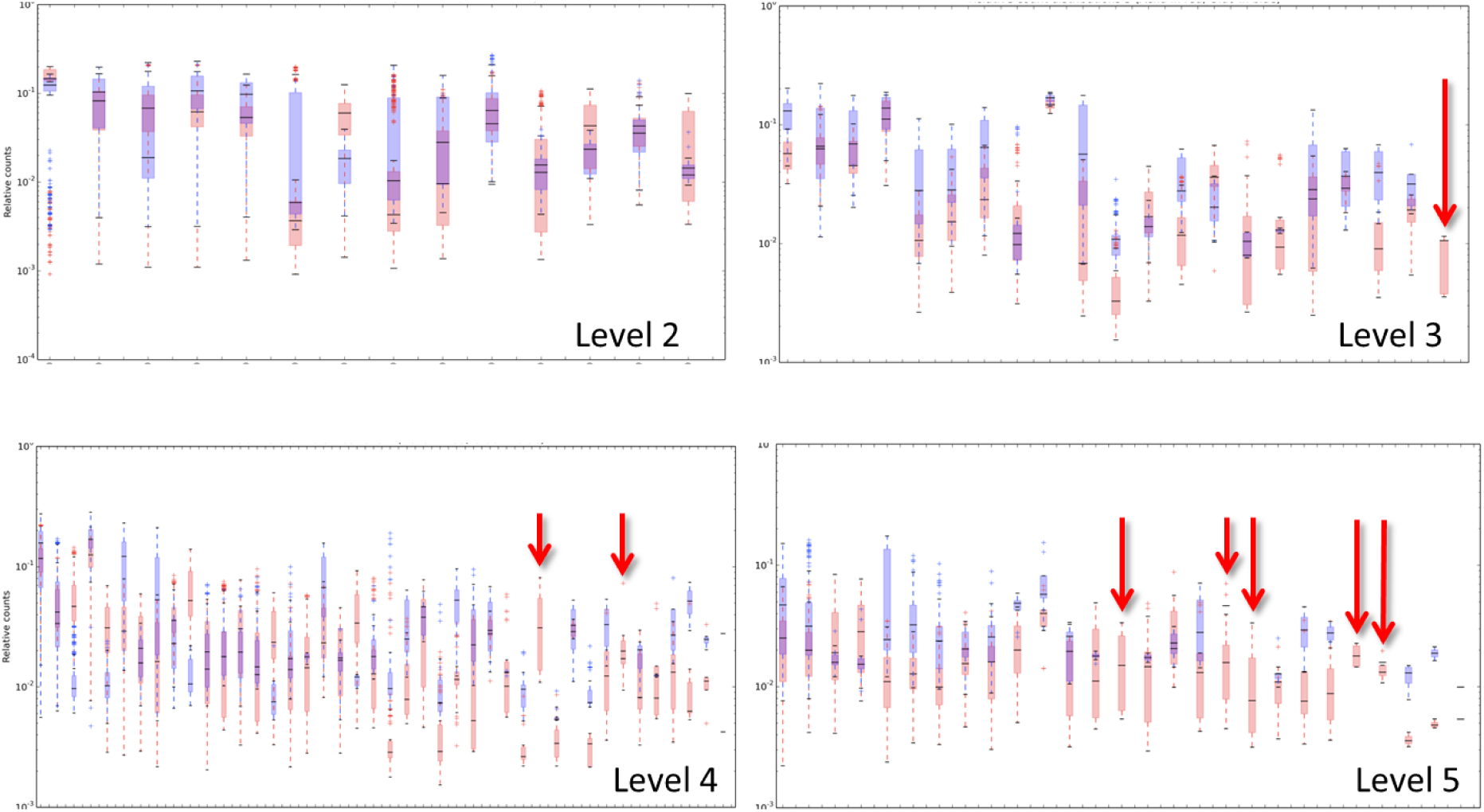
Relative frequency of rule appearance across all Oldowan (blue) and Acheulean (red) sequences. Red arrows indicate rules occurring only in Acheulean sequences.

**Supplementary Figure 3.**
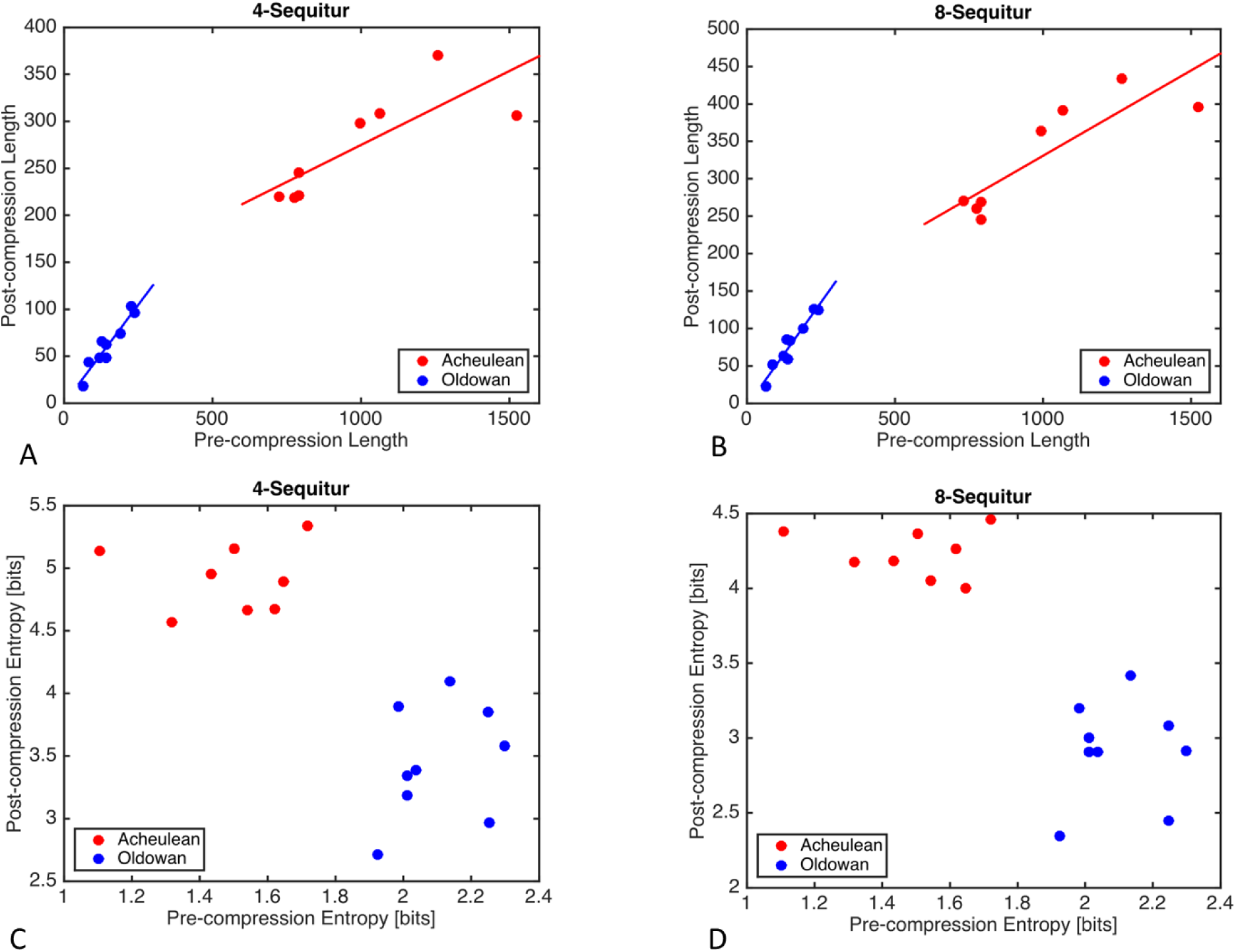
Effect of k-Sequitur compression on sequence length **(A, B)** and entropy **(C,D)** for increasing values of k.

**Supplementary Figure 4.**
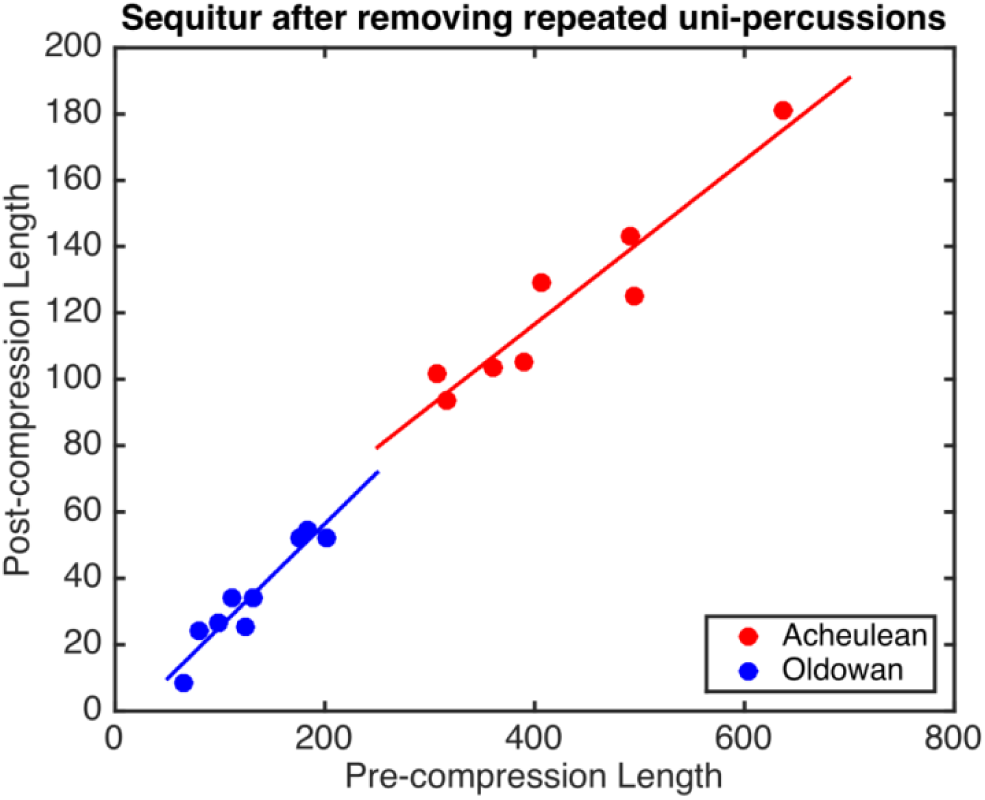
Effect of Sequitur compression after removing repeated percussions.

## Supplementary Discussion: Archaeological Implications

Technological complexity is a fundamental concept in Paleolithic archaeology used to infer everything from cognitive abilities to ecological adaptations and rates of cultural evolution, but has remained poorly defined and impossible to measure in an objective and generalizable way^1^. As a result, interpretations are often driven by qualitative and/or idiosyncratic comparisons between named stone-tool “industries” like the Oldowan and Acheulean that are themselves analytic constructs subsuming a wide range of actual behavioral variation^2^. The Acheulean, for example, spans 1.5 million years of human evolution on three continents. Although its appearance has long been viewed as a major transition in hominin cognitive evolution^3^, more specific links between variation in archaeologically-observable techniques, inferred behavioral complexity, and evolving neurocognitive mechanisms over this vast swath of time and space have remained elusive.

Our current findings focus attention on platform preparation as a key indicator of increasing technological complexity. On an abstract level, platform preparation it is a simple repeat to criterion rule; the difficulty lies in learning to concretely identify the correct criteria for starting and stopping. In other words, it is a problem of learning (through lengthy practice) to perceive subtle properties of the core (e.g. relations between platform convexity, depth, edge angles and core surface morphology^4^) that influence fracture patterns, and then selecting and applying contextually appropriate actions to modify and exploit these properties. To be successful, knapping actions must also conform to very specific kinematic parameters – for example by meeting but not exceeding the kinetic energy required for initiation of the intended fracture^5^. Required energy is in turn a function of the relations between core properties, themselves potentially manipulable through platform preparation, and the size and shape of the desired flake as determined by more distal knapping goals. Thus, the production of effective knapping action sequences requires a tight integration across multiple levels of organization, from the kinematic details of individual blows to the superordinate goals toward which these blows are directed, as well as the ability to flexibly switch between the various sub-goal states that define these relations as the tool-making process unfolds.

The actual Paleolithic use of platform preparation to achieve these ends was confirmed in a study of handaxe production waste from the ˜500,000 year old site of Boxgrove, UK^6^. Platform preparation has also been reported in the context of a different technology (blade production) at the similarly-aged site of Kathu Pan 1 in South Africa^7^. Such research follows well-established experimental archaeology methods for the reconstruction of past behavior by using experimental replication to generate expectations that are testable against actual archaeological materials^8^. To date, however, assessment of the cognitive implications of such reconstructed behaviors has been largely reliant on informal and qualitative interpretation. Neuroimaging studies of experimental tool-making offer the prospect of a more rigorous and empirical approach but face significant challenges of their own, including especially the difficulty of dealing with complex naturalistic behaviors like knapping.

For example, a recent study using functional near-infrared spectroscopy found that rIFG response during experimental handaxe-making was modulated by the presence/absence of linguistic instruction during training^9^, which was manipulated by presenting identical training videos with or without sound. The authors concluded that rIFG activity likely reflected use of inner speech by the linguistically-instructed group rather than the demands of response selection. However rIFG is not one of the cortical regions typically associated with inner speech^10^, and an alternative explanation is that differential training yielded different learning outcomes^11^,^12^ with rIFG responding to overt differences in task performance. Consistent with the latter, the authors do report differences in mean flake shape and relative platform size in debitage produced by the two groups. Properly testing such alternatives will require detailed description and quantification of actual subject behavior. While this is challenging for multi-component, real-world tasks like stone tool-making, grammar extraction using HMM and/or CFG does provide a promising method for data-driven parameterization of the structural complexity hypothesized to drive rIFG response.

## References

1 Fitch, W. & Martins, M. D. Hierarchical processing in music, language, and action: Lashley revisited. Annals of the New York Academy of Sciences 1316, 87–104 (2014).

2 Lashley, K. in Cerebral mechanisms in behavior (*ed* L. A. Jeffress) 112–136 (John Wiley, 1951).

3 Hauser, M. D., Chomsky, N. & Fitch, W. T. The faculty of language: what is it, who has it and how did it evolve? Science 298, 1569–1579 (2002).

4 Byrne, R. & Russon, A. E. Learning by imitation: a hierarchical approach. Behavioral and Brain Sciences 21, 667–721 (1998).

5 Holloway, R. Culture: a human domain. Current Anthropology 10, 395–412 (1969).

6 Greenfield, P. M. Language, tools, and brain: The development and evolution of hierarchically organized sequential behavior. Behavioral and Brain Sciences 14, 531–595 (1991).

7 Botvinick, M. & Weinstein, A. Model-based hierarchical reinforcement learning and human action control. Philosophical Transactions of the Royal Society B: Biological Sciences 369, doi:10.1098/rstb.2013.0480 (2014).

8 Duncan, J. The multiple-demand (MD) system of the primate brain: mental programs for intelligent behaviour. Trends in cognitive sciences 14, 172–179 (2010).

9 Fedorenko, E., Duncan, J. & Kanwisher, N. Language-selective and domain-general regions lie side by side within Broca’s area. Current Biology 22, 2059–2062 (2012).

10 Kolodny, O., Edelman, S. & Lotem, A. Evolution of protolinguistic abilities as a by-product of learning to forage in structured environments. Proc. R. Soc. B 282, 20150353 (2015).

11 Perreault, C., Brantingham, P. J., Kuhn, S. L., Wurz, S. & Gao, X. Measuring the complexity of lithic technology. Current Anthropology 54, S397–S406 (2013).

12 Stout, D. Stone toolmaking and the evolution of human culture and cognition. Philosophical Transactions of the Royal Society B: Biological Sciences 366, 1050–1059 (2011).

13 Mahaney, R. A. Exploring the Complexity and Structure of Acheulean Stoneknapping in Relation to Natural Language. PaleoAnthropology 2014, 586–606 (2014).

14 Faisal, A., Stout, D., Apel, J. & Bradley, B. The Manipulative Complexity of Lower Paleolithic Stone Toolmaking. PLos One 5, e13718 (2010).

15 Stout, D., Apel, J., Commander, J. & Roberts, M. Late Acheulean technology and cognition at Boxgrove, UK. Journal of Archaeological Science 41, 576–590 (2014).

16 Belić, J. J. & Faisal, A. A. Decoding of human hand actions to handle missing limbs in Neuroprosthetics. Frontiers in computational neuroscience 9 (2015).

17 Stout, D., Toth, N., Schick, K. D. & Chaminade, T. Neural correlates of Early Stone Age tool-making: technology, language and cognition in human evolution. Philosophical Transactions of the Royal Society of London B 363, 1939–1949 (2008).

18 Stout, D., Passingham, R., Frith, C., Apel, J. & Chaminade, T. Technology, expertise and social cognition in human evolution. European Journal of Neuroscience 33, 1328–1338, doi:10.1111/j.1460-9568.2011.07619.x (2011).

19 Hecht, E. E. et al. Acquisition of Paleolithic toolmaking abilities involves structural remodeling to inferior frontoparietal regions. Brain Structure and Function, 1–17, doi:10.1007/s00429-014-0789-6 (2014).

20 Engels, F. in Philosophy of Technology (eds R. C. Scharff & Dusek. V.) 71–77 (Blackwell, 2003 [1873]).

21 Ambrose, S. Paleolithic technology and human evolution. Science 291, 1748–1753 (2001).

22 Stout, D. & Hecht, E. E. Evolutionary neuroscience of cumulative culture. Proc Natl Acad Sci U S A, 1–8, doi:www.pnas.org/cgi/doi/10.1073/pnas.1620738114 (in press).

23 Newcomer, M. H. Some Quantitative Experiments in Handaxe Manufacture. World Archaeology 3, 85–104 (1971).

24 Moore, M. W. in Stone tools and the evolution of human cognition (eds April Nowell & Iain Davidson) 13–43 (University Press of Colorado, 2010).

25 Petersson, K.-M., Folia, V. & Hagoort, P. What artificial grammar learning reveals about the neurobiology of syntax. Brain and language 120, 83–95 (2012).

26 Donoso, M., Collins, A. G. E. & Koechlin, E. Foundations of human reasoning in the prefrontal cortex. Science 344, 1481–1486, doi:10.1126/science.1252254 (2014).

27 Frank, S. L., Bod, R. & Christiansen, M. H. How hierarchical is language use? Proceedings of the Royal Society B: Biological Sciences 279, 4522–4531 (2012).

28 Fitch, W. T. & Hauser, M. D. Computational Constraints on Syntactic Processing in a Nonhuman Primate. Science 303, 377–380, doi:10.1126/science.1089401 (2004).

29 Fitch, W. T. Toward a computational framework for cognitive biology: unifying approaches from cognitive neuroscience and comparative cognition. Physics of life reviews 11, 329–364 (2014).

30 Nevill-Manning, C. G. & Witten, I. H. Identifying hierarchical strcture in sequences: A linear-time algorithm. J. Artif. Intell. Res.(JAIR) 7, 67–82 (1997).

31 Schick, K. D. & Toth, N. Making silent stones speak: human evolution and the dawn of technology. (Simon & Schuster, 1993).

32 Putt, S. S., Wijeakumar, S., Franciscus, R. G. & Spencer, J. P. The functional brain networks that underlie Early Stone Age tool manufacture. Nature Human Behaviour 1, 0102 (2017).

33 Aron, A. R., Robbins, T. W. & Poldrack, R. A. Inhibition and the right inferior frontal cortex: one decade on. Trends in cognitive sciences 18, 177–185 (2014).

34 Levy, B. J. & Wagner, A. D. Cognitive control and right ventrolateral prefrontal cortex: reflexive reorienting, motor inhibition, and action updating. Annals of the New York Academy of Sciences 1224, 40–62, doi:10.1111/j.1749-6632.2011.05958.x (2011).

35 Verbruggen, F., Schneider, D. W. & Logan, G. D. How to stop and change a response: the role of goal activation in multitasking. Journal of Experimental Psychology: Human Perception and Performance 34, 1212 (2008).

36 Dippel, G. & Beste, C. A causal role of the right inferior frontal cortex in implementing strategies for multi-component behaviour. Nature communications 6 (2015).

37 Hecht, E. E., Gutman, D. A., Bradley, B. A., Preuss, T. M. & Stout, D. Virtual dissection and comparative connectivity of the superior longitudinal fasciculus in chimpanzees and humans. NeuroImage 108, 124–137, doi:http://dx.doi.org/10.1016/j.neuroimage.2014.12.039 (2015).

38 Stout, D. & Chaminade, T. Stone tools, language and the brain in human evolution. Philosophical Transactions of the Royal Society B: Biological Sciences 367, 75–87, doi:10.1098/rstb.2011.0099 (2012).

39 Stout, D. & Hecht, E. E. Evolutionary neuroscience of cumulative culture. Proceedings of the National Academy of Sciences 114, 7861–7868, doi:10.1073/pnas.1620738114 (2017).

40 Poldrack, Russell A. Inferring Mental States from Neuroimaging Data: From Reverse Inference to Large-Scale Decoding. Neuron 72, 692–697, doi:https://doi.org/10.1016/j.neuron.2011.11.001 (2011).

41 Hutzler, F. Reverse inference is not a fallacy per se: Cognitive processes can be inferred from functional imaging data. Neuroimage 84, 1061–1069 (2014).

42 Stout, D., Hecht, E., Khreisheh, N., Bradley, B. & Chaminade, T. Cognitive Demands of Lower Paleolithic Toolmaking. PLoS ONE 10, e0121804, doi:10.1371/journal.pone.0121804 (2015).

43 Nonaka, T., Bril, B. & Rein, R. How do stone knappers predict and control the outcome of flaking? Implications for understanding early stone tool technology. Journal of Human Evolution 59, 155–167, doi:DOI:10.1016/j.jhevol.2010.04.006 (2010).

44 Stolk, A., Verhagen, L. & Toni, I. Conceptual alignment: How brains achieve mutual understanding. Trends in cognitive sciences 20, 180–191 (2016).

45 Chaminade, T. An experimental approach to study the physiology of natural social interactions. Interaction Studies 18, 254–275, doi:DOI:10.1075/is.18.2.06gry (2017).

46 Brennan, J. Naturalistic Sentence Comprehension in the Brain. Language & Linguistics Compass 10, 299–313, doi:10.1111/lnc3.12198 (2016).

47 Yeo, B. T. et al. The organization of the human cerebral cortex estimated by intrinsic functional connectivity. Journal of neurophysiology 106, 1125–1165 (2011).

48 Keysers, C., Kaas, J. H. & Gazzola, V. Somatosensation in social perception. Nat Rev Neurosci 11, 417–428 (2010).

49 Stout, D. & Chaminade, T. The evolutionary neuroscience of tool making. Neuropsychologia 45, 1091–1100 (2007).

50 Xiloyannis, M., Gavriel, C., Thomik, A. & Faisal, A. Gaussian process autoregression for simultaneous proportional multi-modal prosthetic control with natural hand kinematics.

51 Reznick, R. K. & MacRae, H. Teaching Surgical Skills — Changes in the Wind. New England Journal of Medicine 355, 2664–2669, doi:10.1056/NEJMra054785 (2006).

52 Mnih, V. et al. Human-level control through deep reinforcement learning. Nature 518, 529 (2015).

53 Eren, M. I. et al. Test, Model, and Method Validation: The Role of Experimental Stone Artifact Replication in Hypothesis-driven Archaeology. Ethnoarchaeology 8, 103–136, doi:10.1080/19442890.2016.1213972 (2016).

54 Shea, J. J. Lithic Modes A–I: A New Framework for Describing Global-Scale Variation in Stone Tool Technology Illustrated with Evidence from the East Mediterranean Levant. Journal of Archaeological Method and Theory 20, 151–186 (2013).

55 Toth, N. The Oldowan reassessed: A close look at early stone artifacts. Journal of Archaeological Science 12, 101–120 (1985).

56 Harmand, S. et al. 3.3-million-year-old stone tools from Lomekwi 3, West Turkana, Kenya. Nature 521, 310–315 (2015).

57 Stout, D., Semaw, S., Rogers, M. J. & Cauche, D. Technological variation in the earliest Oldowan from Gona, Afar, Ethiopia. Journal of Human Evolution 58, 474–491, doi:DOI:10.1016/j.jhevol.2010.02.005 (2010).

58 Moore, M. W. The design space of stone flaking: implications for cognitive evolution. World Archaeology 43, 702–715 (2011).

59 Inizan, M.-L., Reduron-Ballinger, M., Roche, H. & Tixier, J. Technology and terminology of knapped stone. (C.R.E.P., 1999).

60 Ottoni, E. B. EthoLog 2.2: a tool for the transcription and timing of behavior observation sessions. Behavior Research Methods, Instruments, & Computers 32, 446–449 (2000).

61 Roux, V. & David, E. in Stone knapping: the necessary conditions for a uniquely hominin behaviour (eds Valentine Roux & Blandine Bril) 91-108 (McDonald Institute for Archaeological Research, 2005).

62 Neumann, N., Lotze, M. & Eickhoff, S. B. Cognitive Expertise: An ALE Meta-Analysis. Human Brain Mapping 37, 262–272, doi:10.1002/hbm.23028 (2016).

63 Jeon, H.-A. & Friederici, A. D. Degree of automaticity and the prefrontal cortex. Trends in cognitive sciences 19, 244–250 (2015).

64 Eickhoff, S. B. et al. Assignment of functional activations to probabilistic cytoarchitectonic areas revisited. Neuroimage 36, 511–521 (2007).

## Supplementary References

1 Perreault, C., Brantingham, P. J., Kuhn, S. L., Wurz, S. & Gao, X. Measuring the complexity of lithic technology. Current Anthropology 54, S397–S406 (2013).

2 Shea, J. J. Lithic Modes A–I: A New Framework for Describing Global-Scale Variation in Stone Tool Technology Illustrated with Evidence from the East Mediterranean Levant. Journal of Archaeological Method and Theory 20, 151–186 (2013).

3 Ambrose, S. Paleolithic technology and human evolution. Science 291, 1748–1753 (2001).

4 Magnani, M., Rezek, Z., Lin, S. C., Chan, A. & Dibble, H. L. Flake variation in relation to the application of force. Journal of Archaeological Science 46, 37–49 (2014).

5 Nonaka, T., Bril, B. & Rein, R. How do stone knappers predict and control the outcome of flaking? Implications for understanding early stone tool technology. Journal of Human Evolution 59, 155–167, doi:DOI:10.1016/j.jhevol.2010.04.006 (2010).

6 Stout, D., Apel, J., Commander, J. & Roberts, M. Late Acheulean technology and cognition at Boxgrove, UK. Journal of Archaeological Science 41, 576–590 (2014).

7 Wilkins, J. & Chazan, M. Blade production~ 500 thousand years ago at Kathu Pan 1, South Africa: support for a multiple origins hypothesis for early Middle Pleistocene blade technologies. Journal of Archaeological Science 39, 1883–1900 (2012).

8 Stout, D. & Khreisheh, N. Skill Learning and Human Brain Evolution: An Experimental Approach. Cambridge Archaeological Journal 25, 867–875, doi:doi:10.1017/S0959774315000359 (2015).

9 Putt, S. S., Wijeakumar, S., Franciscus, R. G. & Spencer, J. P. The functional brain networks that underlie Early Stone Age tool manufacture. Nature Human Behaviour 1, 0102 (2017).

10 Alderson-Day, B. & Fernyhough, C. Inner speech: Development, cognitive functions, phenomenology, and neurobiology. Psychological Bulletin 141, 931–965, doi:10.1037/bul0000021 (2015).

11 Morgan, T. J. et al. Experimental evidence for the co-evolution of hominin tool-making teaching and language. Nature communications 6, 6029, doi:10.1038/ncomms7029 (2015).

12 Putt, S. S., Woods, A. D. & Franciscus, R. G. The Role of Verbal Interaction During Experimental Bifacial Stone Tool Manufacture. Lithic Technology 39, 96–112 (2014).

